# Distinct roles and actions of PDI family enzymes in catalysis of nascent-chain disulfide formation

**DOI:** 10.1101/2021.01.04.425348

**Authors:** Chihiro Hirayama, Kodai Machida, Kentaro Noi, Tadayoshi Murakawa, Masaki Okumura, Teru Ogura, Hiroaki Imataka, Kenji Inaba

## Abstract

The mammalian endoplasmic reticulum (ER) harbors more than 20 members of the protein disulfide isomerase (PDI) family that act to maintain proteostasis. Herein, we developed an *in vitro* system for directly monitoring PDI- or ERp46-catalyzed disulfide bond formation in ribosome-associated nascent chains (RNC) of human serum albumin. The results indicated that ERp46 more efficiently introduced disulfide bonds into nascent chains with short segments exposed outside the ribosome exit site than PDI. Single-molecule analysis by high-speed atomic force microscopy further revealed that PDI binds nascent chains persistently, forming a stable face-to-face homodimer, whereas ERp46 binds for a shorter time in monomeric form, indicating their different mechanisms for substrate recognition and disulfide bond introduction. Similarly to ERp46, a PDI mutant with an occluded substrate-binding pocket displayed shorter-time RNC binding and higher efficiency in disulfide introduction than wild-type PDI. Altogether, ERp46 serves as a more potent disulfide introducer especially during the early stages of translation, whereas PDI can catalyze disulfide formation in RNC when longer nascent chains emerge out from ribosome.

## Introduction

Over billions of years of evolution, living organisms have developed ingenious mechanisms to promote protein folding (Hartl *et al*, 2011). The oxidative network catalyzing protein disulfide bond formation in the endoplasmic reticulum (ER) is a prime example. While canonical protein disulfide isomerase (PDI) and ER oxidoreductin-1 (Ero1) were previously postulated to constitute a primary disulfide bond formation pathway (Araki & Inaba, 2012; Mezghrani *et al*, 2001; Tavender & Bulleid, 2010), more than 20 different PDI family enzymes and multiple PDI oxidases besides Ero1 have recently been identified in the mammalian ER, suggesting the development of highly diverse oxidative networks in higher eukaryotes (Nguyen *et al*, 2011; Schulman *et al*, 2010; Tavender *et al*, 2010). Each PDI family enzyme is likely to play a distinct role in catalyzing the oxidative folding of different substrates, concomitant with some functional redundancy, leading to the efficient production of a wide variety of secretory proteins with multiple disulfide bonds (Bulleid & Ellgaard, 2011; Okumura *et al*, 2015; Sato & Inaba, 2012).

Our previous *in vitro* studies using model substrates such as reduced and denatured bovine pancreatic trypsin inhibitor (BPTI) and ribonuclease A (RNase A) demonstrated that different PDI family enzymes participate in different stages of oxidative protein folding, resulting in the accelerated folding of native enzymes (Kojima *et al*, 2014; Sato *et al*, 2013). Multiple PDI family enzymes cooperate to synergistically increase the speed and fidelity of disulfide bond formation in substrate proteins. However, whether mechanistic insights gained by *in vitro* experiments using full-length substrates are applicable to real events of oxidative folding in the ER remains an important question. Indeed, some previous works demonstrated that newly synthesized polypeptide chains undergo disulfide bond formation and isomerization co-translationally, presumably via catalysis by specific PDI family members (Kadokura *et al*, 2020; Molinari & Helenius, 1999; Robinson & Bulleid, 2020; Robinson *et al*, 2020; Robinson *et al*, 2017). Furthermore, nascent chains play important roles in their own quality control by modulating the translation speed to increase the yield of native folding; if a nascent chain fails to fold or complete translation, then the resultant aberrant ribosome-nascent chain complexes are degraded or destabilized (Buhr *et al*, 2016; Chadani *et al*, 2017; Matsuo *et al*, 2017). These observations suggest that understanding real events of oxidative protein folding in cells requires systematic analysis of how PDI family enzymes act on nascent polypeptide chains during synthesis by ribosomes.

To this end, we herein developed an experimental system for directly monitoring disulfide bond formation in ribosome-associated human serum albumin (HSA) nascent chains of different lengths from the N-terminus. The resultant ribosome-nascent chain complexes (RNCs) were reacted with two ubiquitously expressed PDI family members, ER-resident protein 46 (ERp46) and canonical PDI. These two enzymes were previously shown to have distinct roles in catalyzing oxidative protein folding: ERp46 engages in rapid but promiscuous disulfide bond introduction during the early stages of folding, while PDI serves as an effective proofreader of non-native disulfides during the later stages (Kojima *et al*., 2014; Sato *et al*., 2013). The subsequent maleimidyl polyethylene glycol (mal-PEG) modification of free cysteines and Bis-Tris (pH7.0) PAGE analysis enabled us to detect the oxidation status of the HSA nascent chains conjugated with transfer RNA (tRNA). Using high-speed atomic force microscopy (HS-AFM), we further visualized PDI and ERp46 acting on the RNCs at the single-molecule level. Collectively, the results indicated that although both ERp46 and PDI could introduce a disulfide bond into the ribosome-associated HSA nascent chains, they demanded different lengths of the HSA segment exposed outside the ribosome exit site, and displayed different mechanisms of action against the RNC. The present systematic *in vitro* study using RNC containing different lengths of HSA nascent chains mimics co-translational disulfide bond formation in the ER, and the results provide a framework for understanding the mechanistic basis of oxidative nascent-chain folding catalyzed by PDI family enzymes.

## Results

### The efficiency of disulfide bond introduction into HSA nascent chains by PDI/ERp46

To investigate whether PDI family enzymes can introduce disulfide bonds into a substrate during translation, we first prepared RNCs *in vitro*. For this purpose, we made use of a cell-free protein translation system reconstituted with eukaryotic elongation factors 1 and 2, eukaryotic release factors 1 and 3 (eRF1 and eRF3), aminoacyl-tRNA synthetases, tRNAs, and ribosome subunits, developed previously by Imataka and colleagues (Machida *et al*, 2014). HSA was chosen as a model substrate for the following reasons. Firstly, the three-dimensional structure of HSA has been solved at high resolution (Sugio *et al*, 1999), providing information on the exact location of 17 disulfide bonds in its native structure. Secondly, native-state HSA contains an unpaired cysteine, Cys34, near the N-terminal region, which has potential to form a non-native disulfide bond with one of the subsequent cysteines, serving as a good indicator of whether a non-native disulfide is introduced by ERp46 or PDI during the early stage of translation. Thirdly, overall conformation and kinetics of disulfide bond regeneration were characterized for reduced full-length HSA (Lee & Hirose, 1992), which is beneficial for discussing similarities and differences in post- and co-translational oxidative folding. Forth, no N-glycosylation sites are contained in the first 95 amino acids of HSA, implying that HSA nascent chains synthesized by the cell-free system are equivalent to those synthesized in the ER in regard to N-glycosylation. Finally, the involvement of PDI family enzymes in intracellular HSA folding has been demonstrated (Koritzinsky *et al*, 2013; Rutkevich *et al*, 2010; Rutkevich & Williams, 2012), ensuring the physiological relevance of the present study.

To stall the translation of HSA at specified sites, a uORF2 arrest sequence (Alderete *et al*, 1999) was inserted into appropriate sites of the expression plasmid (Fig 1A). We first prepared two versions of the RNC containing different lengths of HSA nascent chains: RNC 69-aa and RNC 82-aa. Since the ribosome exit tunnel accommodates a polypeptide chain of ∼30 amino acid (aa) residues (Zhang *et al*, 2013), the N-terminal 57 residues of HSA (excluding the N-terminal 6-aa pro-sequence) are predicted to be exposed outside the ribosome exit tunnel in RNC 69-aa, including Cys34 and Cys53 (Fig 1A). In the RNC 82-aa construct, the N-terminal 70 residues of HSA, including Cys62 as well as Cys34/Cys53, are predicted to emerge from the ribosome (Fig 1A). Notably, Cys53 and Cys62 form a native disulfide bond, whereas Cys34 is unpaired in the native structure of HSA domain I.

**Figure 1.**
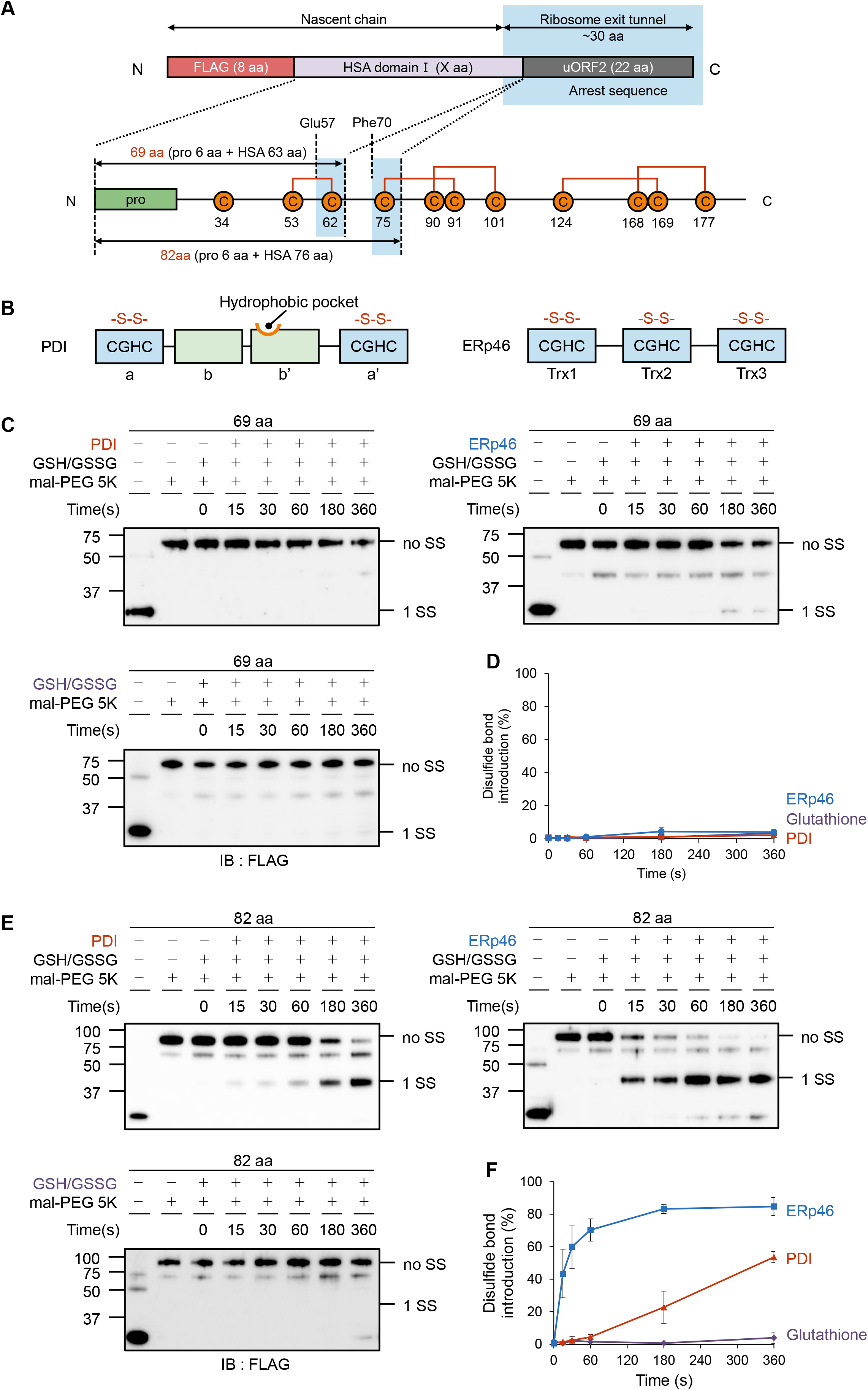
Disulfide bond introduction into RNC 69-aa and 82-aa by PDI and ERp46. **A** Schematic structure of plasmids constructed in this study. ‘uORF2’ is an arrest sequence that serves to stall translation of the upstream protein and thereby prepare stable ribosome-nascent chain complexes (RNCs). The bottom cartoon represents the location of cysteines and disulfide bonds in HSA domain I. HSA domain I consists of 195 amino acids and contains five disulfide bonds and one free cysteine at residue 34. A green box indicates the pro-sequence. Orange circles and red lines indicate cysteines and native disulfide bonds, respectively. The region predicted to be buried in the ribosome exit tunnel is shown by a cyan box. **B** Domain organization of PDI and ERp46. Redox-active Trx-like domains with a CGHC motif are indicated by cyan boxes, while redox-inactive ones in PDI are by light-green boxes. Note that the PDI **b’** domain contains a substrate-binding hydrophobic pocket. **C, E** Time course of PDI-, ERp46-, and glutathione (no enzyme)-catalyzed disulfide bond introduction into RNC 69-aa (C) and 82-aa (E). ‘noSS’ and ‘1SS’ denote reduced and single-disulfide-bonded species of HSA nascent chains, respectively. Note that faint bands observed between “no SS” and “1SS” likely represent a species in which one of cysteines is not subjected to mal-PEG modification due to glutathionylation. In support of this, these minor bands are even fainter under the conditions of no GSH/GSSG. **D, F** Quantification of disulfide-bonded species for RNC 69-aa (D) and 82-aa (F) based on the results shown in (C) and (E), respectively (n = 3).

When RNC 69-aa was employed as a substrate, neither PDI nor ERp46 could efficiently introduce a disulfide bond into the nascent chain (Fig 1C and 1D). However, both enzymes introduced a disulfide bond into RNC 82-aa with higher efficiency than into RNC 69-aa (Fig 1E and 1F), suggesting that the length of the exposed HAS segment or the distance of a pair of cysteines from the ribosome exit site is critical for disulfide bond introduction by PDI and ERp46. For either construct, a faint band was seen between the bands of ‘no SS’ and ‘1 SS’, and this band was even fainter without GSH/GSSG (the second lane from the left) and had a tendency to get stronger at late time points. Presumably, this band represents a species in which one of free cysteines is glutathionylated, and the species increased gradually in the course of the reaction.

Of note, ERp46 introduced a disulfide bond into RNC 82-aa at a much higher rate than PDI, indicating that ERp46 serves as a more competent disulfide bond introducer to RNCs than PDI (Fig 1F). The remarkable difference in disulfide bond introduction efficiency by these two enzymes seems unlikely to be explained simply by the different number of redox-active Trx-like domains in PDI (two) and ERp46 (three) (Fig 1B). Also, the redox states in the presence of 1 mM GSH and 0.2 mM GSSG are similar between these two enzymes (Fig EV1A and EV1B), suggesting their comparable redox potentials. Thus, the different ability of ERp46 and PDI to introduce a disulfide into 82-aa is likely caused by other factors such as different structural features and different mechanism of substrate recognition, as discussed below.

Next, to identify which cysteine pair forms a disulfide bond in RNC 82-aa, we constructed three cysteine mutants in which either Cys34, Cys53, or Cys62 was mutated to alanine (Fig 2A). The assays using the mutants showed that whereas PDI was unable to introduce a disulfide bond into RNC 82-aa C34A and C53A (Fig 2B, top and middle), the enzyme introduced a Cys34-Cys53 non-native disulfide bond into RNC 82-aa C62A (Fig 2B, bottom), at almost the same rate as the generation of the ‘1 SS’ species in 82-aa (Fig 1E and 1F). PDI could not introduce a Cys53-Cys62 native disulfide bond, presumably because this cysteine pair is located too close to the ribosome exit site (see also Fig 3B and 3C). Conversely, the slow but possible formation of a Cys34-Cys53 non-native disulfide in 82 aa by PDI suggests that the distance between a cysteine pair of interest and the ribosome exit site is key to allowing the enzyme to catalyze disulfide bond introduction into RNCs. Considering the different locations of the Cys34-Cys53 and Cys53-Cys62 pairs on RNC 82-aa, a distance of ∼18 residues from the ribosome exit site appears to be necessary for the PDI-catalyzed reaction (see also the Discussion).

**Figure 2.**
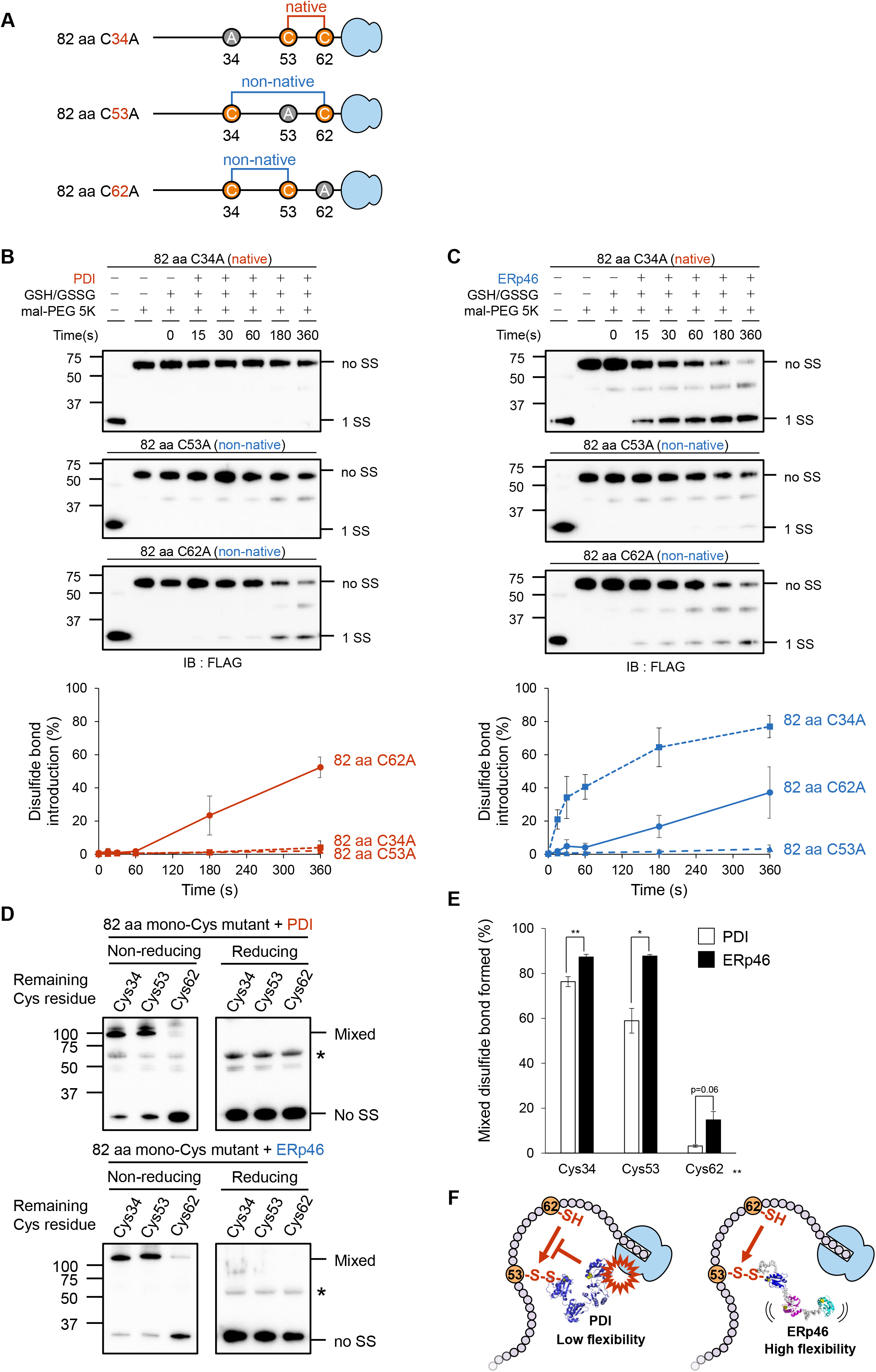
Disulfide bond introduction into RNC 82-aa Cys mutants by PDI and ERp46. **A** Cartoon of RNC constructs used in this study. In each construct, a cysteine (represented by a black circle) was mutated to alanine. Note that RNC 82-aa C34A retains a native cysteine pairing (i.e., Cys53 and Cys62), while RNC 82-aa C53A and C62A retain a non-native pairing. **B** and **C** Time course of PDI- and ERp46-catalyzed disulfide bond introduction into RNC 82-aa C34A (top), C53A (middle), and C62A (bottom) mutants. Note that faint bands observed between “no SS” and “1SS” likely represent a species in which one of cysteines is not subjected to mal-PEG modification due to glutathionylation. Quantification of disulfide-bonded species of RNC 82-aa Cys mutants is based on the results shown for the upper raw data (n = 3). **D** Formation of a mixed disulfide bond between RNC 82-aa mono-Cys mutants and PDI (upper)/ERp46 (lower). ‘Mixed’ and ‘No SS’ denote a mixed disulfide complex between PDI/ERp46 and RNC mono-Cys mutants and isolated RNC 82-aa, respectively. Note that faint bands observed between ‘Mixed’ and ‘no SS’ are likely non-specific bands, as they were seen at the same position regardless of which 82-aa mono-Cys mutant was tested or whether an RNC was reacted with PDI or ERp46. **E** Quantification of mixed disulfide species based on the results shown in (D). n = 3. **F** The cartoon on the left shows possible steric collisions between ribosomes and PDI when Cys62 attacks the mixed disulfide between Cys53 on RNC 82-aa and PDI (left). The cartoon on the right shows that ERp46 can avoid this steric collision due to its higher flexibility and domain arrangement.

**Figure 3.**
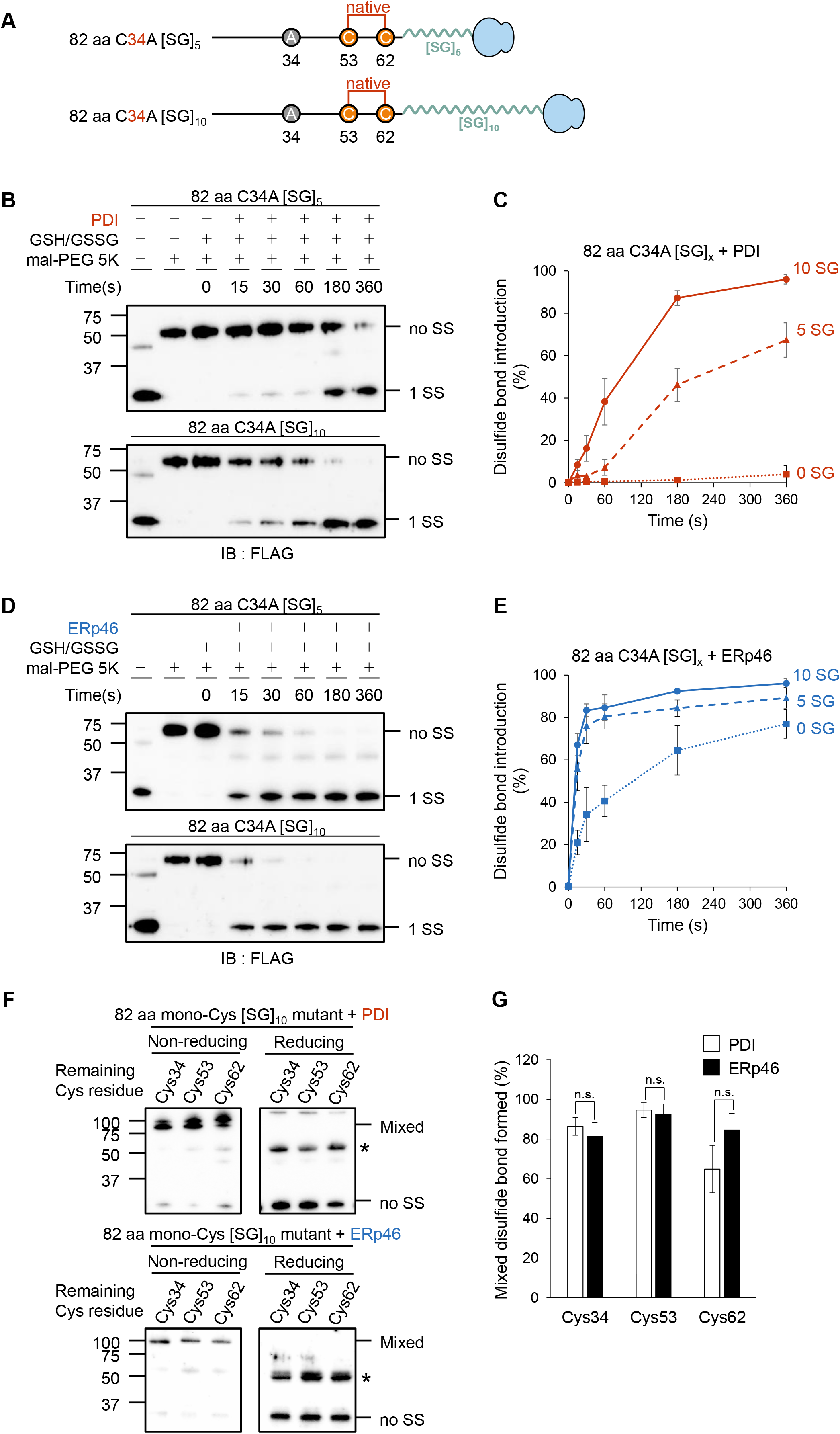
Correlation of the distance between Cys residues and the ribosome exit site with the efficiency of disulfide bond introduction by PDI/ERp46. **A** Cartoons of RNC constructs with [SG]-repeat insertions. A [SG]_5_ or [SG]_10_ repeat sequence was inserted into RNC-82 aa C34A immediately after Cys62. **B, D** PDI-(B) and ERp46 (D)-mediated disulfide bond introduction into RNC 82-aa C34A with insertion of [SG]_5_ (upper) or [SG]_10_ (lower) repeats after Cys62. **C, E** Quantification of disulfide-bonded species (1SS) based on the results shown in (B) and (D). n = 3 for PDI and 2 for ERp46. **F** Formation of a mixed disulfide bond between the 82-aa mono-Cys mutant with a [SG]_10_ repeat and PDI (upper)/ERp46 (lower). Note that bands observed between ‘Mixed’ and ‘no SS’ are likely non-specific bands, as they were seen at the same position regardless of which 82-aa mono-Cys [SG]_10_ mutant was tested or whether an RNC was reacted with PDI or ERp46. **G** Quantification of mixed disulfide species based on the results shown in (F). n = 3.

In contrast to PDI, ERp46 could introduce a native disulfide bond into RNC 82-aa C34A (Fig 2C, top). Like PDI, ERp46 also introduced a non-native disulfide bond between Cys34 and Cys53 into RNC 82-aa C62A, but its efficiency was lower than that of a Cys53-Cys62 native disulfide (Fig 2C, bottom). No disulfide bond was formed between Cys34 and Cys62 by either ERp46 or PDI (Fig 2C, middle), presumably due to the considerable spatial separation of these two cysteines. Based on these results, we concluded that for efficient disulfide bond introduction into RNCs, ERp46 requires an intermediary polypeptide segment with a shorter distance between a cysteine pair of interest and the ribosome exit site than PDI. We here note that ERp46-catalyzed generation of the ‘1 SS’ species was faster in 82-aa than in 82-aa C34A (Fig 1F and 2C). This observation may suggest the occurrence of Cys34-mediated disulfide bond formation in 82-aa, namely, the formation of a Cys34-Cys53 non-native disulfide and, possibly, its rapid isomerization to a Cys53-Cys62 native disulfide.

### Accessibility of PDI/ERp46 to cysteines on the ribosome-HSA nascent chain complex

To examine the accessibility of PDI and ERp46 to Cys residues on RNC 82-aa, we constructed three RNC 82-aa mono-Cys mutants in which either Cys34, Cys53, or Cys62 on the HSA nascent chain was retained, and investigated whether a mixed disulfide could be formed between the RNC 82-aa mutant and a trapping mutant of PDI or ERp46 in which all CXXC redox-active sites were mutated to CXXA. Both PDI and ERp46 formed a mixed disulfide bond with Cys34 and Cys53 on RNC 82-aa with high probability, but covalent linkages to Cys62 were marginal (Fig 2D and 2E). The results suggest that the redox-active sites of PDI and ERp46 could gain access to Cys34 and Cys53, but to a much lesser extent, to Cys62, probably due to steric collision with the ribosome. Nevertheless, ERp46 efficiently introduced a native disulfide bond between Cys53 and Cys62 (Fig 2C, top), presumably because ERp46 first attacked Cys53 on the HSA nascent chain, and the resultant mixed disulfide was subjected to nucleophilic attack by Cys62 (Fig 2F, right). By contrast, the mixed disulfide between PDI and Cys53 on the HSA nascent chain seems unlikely to be attacked by Cys62, probably due to steric collision between PDI and the ribosome (Fig 2F, left). In line with this idea, PDI adopts a U-like overall conformation with restricted movements of four thioredoxin (Trx)-like domains (Tian *et al*, 2006; Wang *et al*, 2012), whereas ERp46 forms a highly flexible V-shape conformation composed of three Trx-like domains and two long (∼20 aa) interdomain linkers (Kojima *et al*., 2014).

### Correlations between cysteine accessibility and the efficiency of disulfide bond introduction by PDI/ERp46

Based on the results presented above, we believe that the distance between cysteines of interest and the ribosome exit site is critical for efficient disulfide introduction by PDI and ERp46. To test this hypothesis, we increased the distance of the Cys53-Cys62 pair from the ribosome exit site by inserting an extended polypeptide segment composed of [SG]_5_ or [SG]_10_ repeat immediately after Cys62 on RNC 82-aa C34A (Fig 3A), and investigated the effects of the insertions on the efficiency of disulfide bond formation. While PDI was unable to introduce a Cys53-Cys62 native disulfide into RNC 82-aa C34A (Fig 2B, top), insertion of a [SG]_5_ repeat allowed this reaction, and nearly 70% of 82-aa C34A was disulfide-bonded within a reaction time of 360 s (Fig 3B, upper and 3C). The insertion of a longer repeat [SG]_10_ further promoted disulfide bond formation (Fig 3B, lower and 3C).

A similar enhancement following [SG] repeat insertion was observed for ERp46-catalyzed reactions. However, ERp46 exhibited a striking difference from PDI: insertion of a [SG]_5_ repeat was long enough to introduce a Cys53-Cys62 native disulfide into RNC 82-aa C34A within 15 s, and insertion of a [SG]_10_ repeat gave only a small additional enhancement (Fig 3D and 3E). Thus, the presence of a disordered or extended segment of ∼18 aa (Asp63−Phe70 + [SG]_5_ repeat) between a cysteine pair of interest and the ribosome exit site was necessary and sufficient for ERp46 to generate a Cys53-Cys62 disulfide rapidly, whereas PDI required a longer segment of ∼28 aa (Asp63−Phe70 + [SG]_10_ repeat) in this intermediary region for efficient introduction of a Cys53-Cys62 disulfide. Thus, ERp46 seems to be more capable of introducing a disulfide bond near the ribosome exit site than PDI. In other words, ERp46 likely has the higher potential to introduce a disulfide bond into the HSA nascent chain during the earlier stages of translation than PDI.

To verify that Cys53-Cys62 disulfide formation facilitated by [SG]_10_ repeat insertion was ascribed to higher accessibility of PDI/ERp46 to Cys62, we again investigated mixed disulfide bond formation between trapping mutants of PDI/ERp46 and each cysteine on RNC 82-aa following [SG]_10_ repeat insertion. Both PDI and ERp46 formed a mixed disulfide with all cysteines including Cys62 (Fig 3F and 3G), indicating that there is a correlation between the accessibility of PDI/ERp46 to a target pair of cysteines and the efficiency of disulfide bond introduction by the enzymes.

### Disulfide bond introduction into a longer HSA nascent chain by PDI/ERp46

In addition to the [SG]-repeat insertion, we examined the effect of natural HSA sequence extension on PDI- or ERp46-mediated disulfide formation. For this purpose, we prepared RNC 95-aa in which the N-terminal 83 amino acids of HSA (excluding the N-terminal 6-aa pro-sequence), including Cys34, Cys53, Cys62, and Cys75, are predicted to emerge from ribosome (Fig 4A). With this construct, however, we had a technical problem with detection of the reduced species, because mal-PEG modification of four cysteines greatly diminished the gel-to-membrane transfer efficiency. We overcame this problem by using photo-cleavable mal-PEG (PEG-PCMal) and irradiating UV light to the SDS gel after the gel electrophoresis and before the membrane transfer.

**Figure 4.**
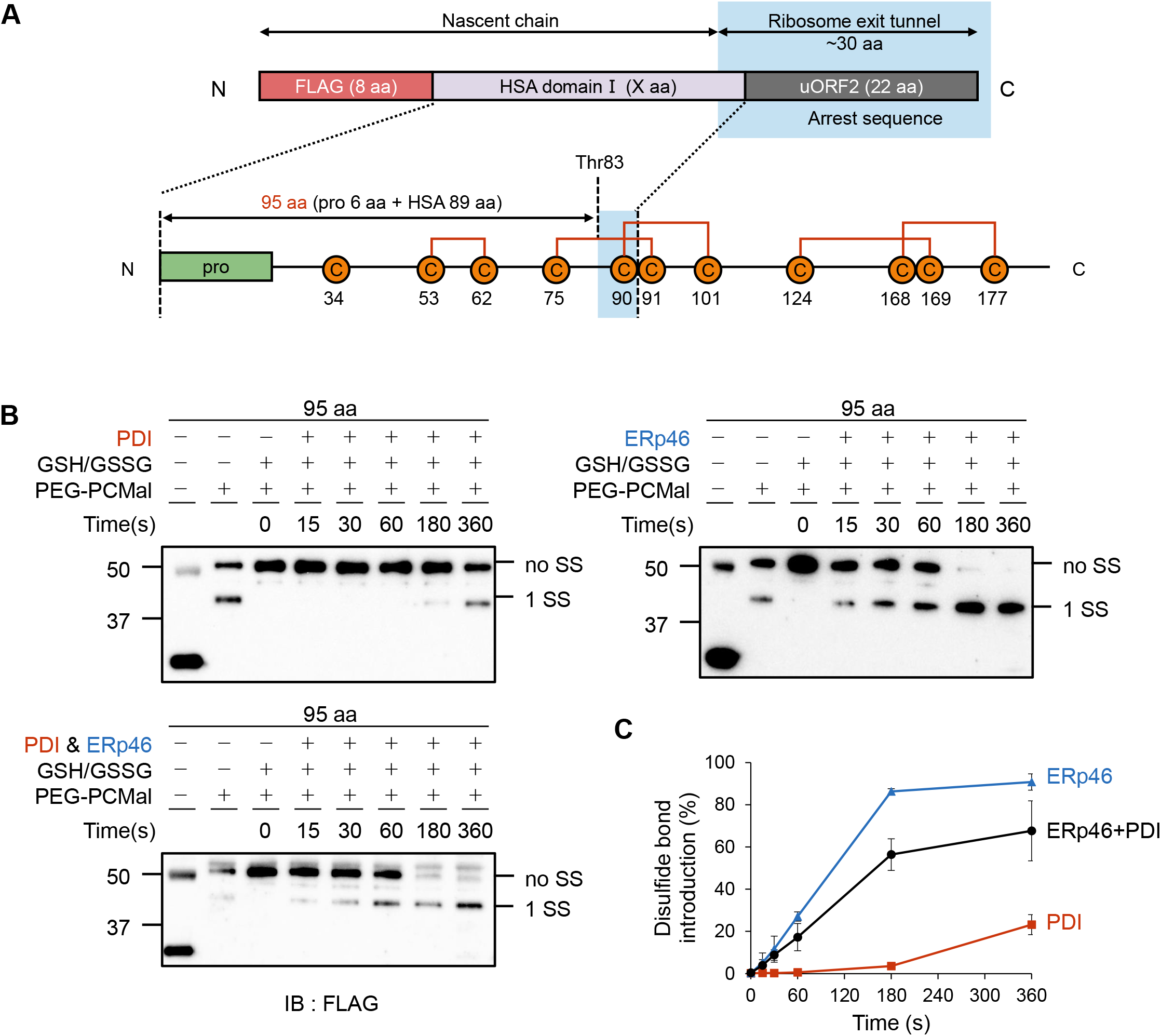
Disulfide bond introduction into RNC 95-aa by PDI and ERp46. **A** Schematic structure of RNC-95-aa. Orange circles and red lines in the bottom cartoon indicate cysteines and native disulfides, respectively. The region predicted to be buried in the ribosome exit tunnel is shown by a cyan box. **B** Time course of PDI (0.1 μM)-, ERp46 (0.1 μM)-, and their mixture (0.1 μM each)-catalyzed disulfide bond introduction into RNC 95-aa. ‘noSS’ and ‘1SS’ denote reduced and single-disulfide-bonded species of the HSA nascent chain, respectively. **C** Quantification of the single-disulfide-bonded (1 SS) species based on the result shown in (B) (n = 3).

Consequently, we observed both PDI and ERp46 introduced a disulfide bond into 95-aa (Fig 4B), but the efficiency was slower than that into 82-aa (Fig 1E and 1F), although a longer polypeptide chain is exposed outside the ribosome exit site in RNC 95-aa. Thus, the effect of natural sequence extension was opposite to that of [SG]-repeat insertion. Formation of some higher-order structure or exposure of another cysteine may somehow prevent PDI and ERp46 from introducing a disulfide bond into RNC 95-aa. Thus, a longer polypeptide chain exposed outside ribosome does not always lead to a higher disulfide formation rate. Rather, it is suggested that PDI and ERp46 can introduce a disulfide bond into a nascent chain with higher efficiency when the necessary and minimum length emerges out.

Given that four cysteines are exposed outside the ribosome in RNC 95-aa, we next investigated whether PDI and ERp46 can catalyze nascent-chain disulfide formation additionally or synergistically. The mixture of PDI and ERp46 generated a ‘1 SS’ species, but not a ‘2 SS’ species, like PDI or ERp46 alone (Fig 4B and 4C). Notably, the presence of PDI inhibited ERp46-mediated disulfide formation, possibly due to its competition with ERp46 for binding to RNC 95-aa. Thus, neither additional nor synergistic effect was observed (Fig 4B and 4C). In this regard, our previous observation for the synergistic cooperation of PDI and ERp46 in RNase A oxidative folding (Sato *et al*., 2013) was not true for the ribosome-associated HSA nascent chain.

### Single-molecule analysis of ERp46 by high-speed atomic force microscopy

To explore the mechanisms by which PDI and ERp46 recognize and act on RNCs at the molecular level, we employed HS-AFM (Kodera *et al*, 2010; Noi *et al*, 2013; Okumura *et al*, 2019; Uchihashi *et al*, 2018). While our previous HS-AFM analysis revealed that PDI molecules form homodimers in the presence of unfolded substrates (Okumura *et al*., 2019), the structure and dynamics of ERp46 have not been analyzed using this experimental approach. Therefore, we first observed ERp46 molecules alone by immobilizing the N-terminal His-tag on a Co^2+^-coated mica surface. AFM images revealed various overall shapes of ERp46 (Fig 5A), and some particle images clearly demonstrated the presence of three thioredoxin (Trx)-like domains in ERp46 (Fig 5A, left). To assess the overall structures of ERp46, we calculated the circularity of each molecule and performed statistical analysis (Uchihashi *et al*., 2018). Circularity is a measure of how circular the outline of an observed molecule is, defined by the equation 4πS/L^2^, where L and S are the contour length of the outline and the area surrounded by the outline, respectively. Thus, a circularity of 1.0 indicates a perfect circle, and values <1 indicate a more extended conformation.

**Figure 5.**
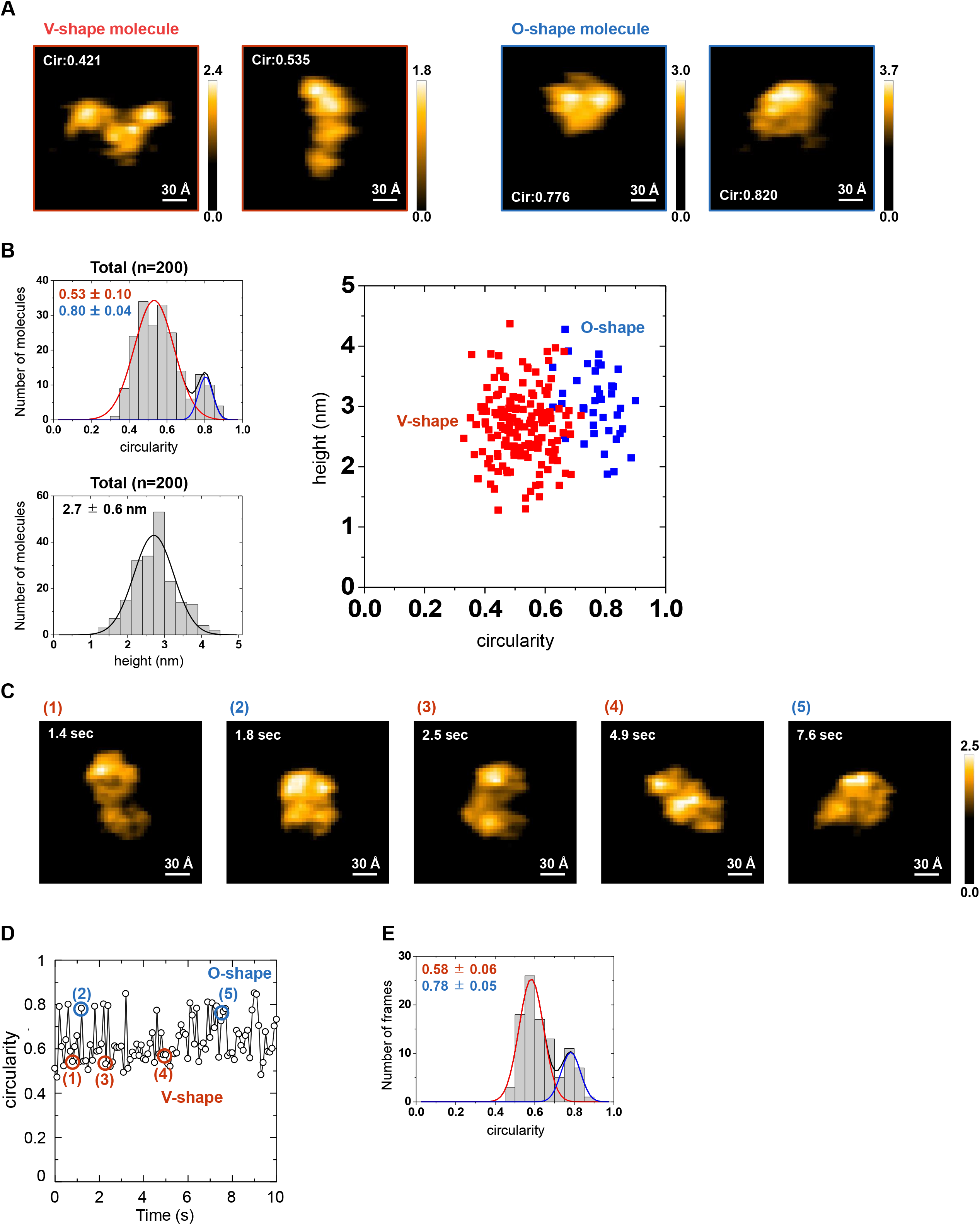
High-speed AFM analysis of ERp46. **A** AFM images (scan area, 200 ’ 200 Å; scale bar, 30 Å) for ERp46 V-shape (left) and O-shape (right) conformations. **B** Left upper: Histograms of circularity calculated from AFM images of ERp46. Values represent the average circularity (mean ± s.d.) calculated from curve fitting with a single-(middle and right) or two-(left) Gaussian model. Left lower: Histograms of height calculated from AFM images of ERp46. Values represent the average height (mean ± s.d.) calculated from curve fitting with a single-Gaussian model. Right: Two-dimensional scatterplots of the height versus circularity for ERp46 molecules observed by HS-AFM. **C** Time-course snapshots of oxidized ERp46 captured by HS-AFM. The images were traced for 10 s. See also Movie EV1. **D** Time trace of the circularity of an ERp46 molecule. **E** Histogram of the circularity of ERp46 calculated from the time-course snapshots shown in (D).

Statistical analysis based on circularity classified randomly chosen ERp46 particles into two major groups: opened V-shape and round/compact O-shape (Fig 5A). Histograms with Gaussian fitting curves indicated that ∼80% of ERp46 molecules adopted V-shape conformations while ∼20% adopted O-shape conformations (Fig 5B). There was no large difference in height between these two conformations, suggesting that the three Trx-like domains of ERp46 are arranged within the same plane in either conformation. Successive AFM images acquired every 100 ms revealed that ERp46 adopted an open V-shape conformation during nearly 75% of the observation time, while the protein also adopted an O-shape conformation occasionally (Fig 5C, 5D, 5E and Movie EV1). The histogram calculated from the time-course snapshots was similar to that calculated from images of 200 molecules at a certain timepoint (Fig 5B and 5E). Importantly, structural insights gained by HS-AFM analysis are in good agreement with those from small-angle X-ray scattering (SAXS) analysis: both analyses consistently indicate the coexistence of a major population of molecules with an open V-shape and a minor population with a compact O-shape (Kojima *et al*., 2014).

### Single-molecule analysis of PDI/ERp46 acting on 82-aa RNC by HS-AFM

PDI and ERp46 are predicted to bind RNCs transiently during disulfide bond introduction, but transient interactions would make it harder to observe and analyze the mode of PDI/ERp46 binding to RNCs. More practically, at least 5 mins are required to prepare for starting HS-AFM measurements after adding PDI or ERp46 to RNCs immobilized onto a mica surface. If we employed RNCs containing natural HSA sequences, PDI or ERp46 would complete nascent-chain disulfide formation during this setup time. We therefore constructed HSA 82-aa RNC with Cys34, Cys53, and Cys62 mutated to Ala (hereafter referred to as 82-aa CA RNC), with the intension of trapping RNC molecules bound to PDI/ERp46. After testing several RNC immobilization methods, we chose to immobilize RNC on a Ni^2+^-coated mica surface. As a result, most RNC molecules were observed to lie sideways on the mica surface, while nascent chains were difficult to visualize, probably due to their flexible and extended structural nature (Fig 6A).

**Figure 6.**
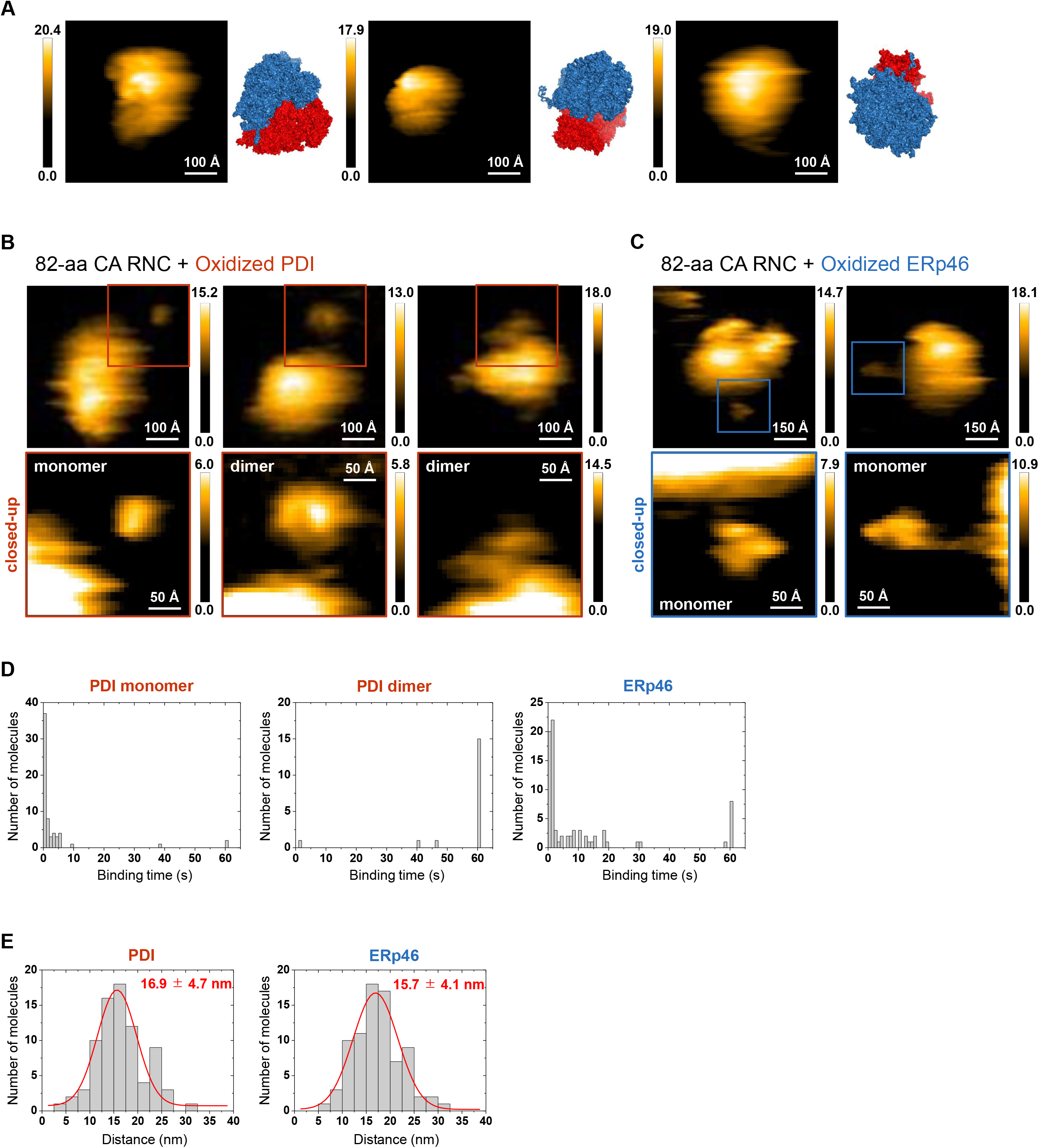
Single-molecule observation of PDI/ERp46 acting on 82-aa CA RNC by high-speed atomic force microscopy. **A** The AFM images (scan area, 500 Å × 500 Å; scale bar, 100 Å) displaying 82-aa CA RNC in the absence of PDI family enzymes on a Ni^2+^-coated mica surface. The surface model on the right side of each AFM image illustrates ribosome whose view angle is approximately adjusted to the observed RNC particle. 40S and 60S ribosomal subunits are shown in red and blue, respectively. **B** Upper AFM images (scan area, 500 Å × 500 Å; scale bar, 100 Å) displaying 82-aa CA RNC in the presence of oxidized PDI (1 nM). PDI molecules that appear to bind 82-aa CA RNC are marked by red squares. Lower images (scan area, 250 Å × 250 Å; scale bar, 50 Å) highlight the regions surrounded by red squares in the upper images. **C** Upper AFM images (scan area, 500 Å × 500 Å; scale bar, 100 Å) displaying 82-aa CA RNC in the presence of oxidized ERp46 (1 nM). ERp46 molecules that appear to bind 82-aa CA RNC are marked by blue squares. Lower images (scan area, 250 Å × 250 Å; scale bar, 50 Å) highlight the regions surrounded by blue squares in the upper images. **D** Histograms of the RNC binding time of the PDI monomer (left), the PDI dimer (middle), and ERp46 (right), calculated from the observed AFM images. **E** Histograms of the distance between the edge of the ribosome and the centers of RNC-neighboring PDI (left) and ERp46 (right) molecules, calculated from the observed AFM images. Values represent the average distance (mean ± s.d.) calculated from curve fitting with a single-Gaussian model.

When oxidized PDI or ERp46 were added to onto the RNC-immobilized mica surface, PDI/ERp46-like particles were observed in the peripheral region of ribosomes. When no-chain RNC (NC-RNC), comprising only the N-terminal FLAG tag and the subsequent uORF2 but no segment from HSA, was immobilized on the mica surface, far fewer particles were observed near RNCs (within 25 Å from the outline of ribosomes) by HS-AFM despite the presence of PDI/ERp46 (Fig EV2A and EV2B). These results confirm that we successfully observed PDI/ERp46 molecules acting on HSA nascent chains associated with ribosomes.

Notably, the HS-AFM analysis revealed that PDI bound RNCs in both monomeric and dimeric forms at an approximate ratio of 7:3 (Fig 6B), as reported previously for reduced and denatured BPTI and RNase A as substrates (Okumura *et al*., 2019). Thus, PDI likely recognizes HSA nascent chains in a similar manner to full-length substrates. Statistical analysis of RNC binding rates revealed that whereas most monomeric PDI molecules (52/55 molecules) bound RNC for 10 s or shorter (Fig 6D, Fig EV3A and Movie EV2), most homodimeric PDI molecules (17/19 molecules) bound RNC for 60 s or longer (Fig 6D, Fig EV3B and Movie EV3). By contrast, ERp46 molecules in the periphery of RNCs were only present in monomeric form (Fig 6C). Importantly, nearly 20% (12/59 molecules) of ERp46 molecules bound RNC for 10 to 20 s (Fig 6D, Fig EV3C and Movie EV4), while a smaller portion (8/59 molecules) bound RNC for ∼60 s (Fig 6D). It is also notable that significant portion of PDI and ERp46 molecules bound ribosomes for <5 s. This may indicate that PDI/ERp46 binds or approaches RNCs only transiently possibly via diffusion, without tight interactions.

The histogram of the distance between the edge of ribosomes and the center of ribosome-neighboring PDI/ERp46 molecules indicated that both PDI and ERp46 bound RNCs at positions ∼16 nm distant from ribosomes with a single-Gaussian distribution with a half width of ∼11 nm (Fig 6E), suggesting that both enzymes recognize similar sites of the HSA nascent chain. Given that the distance between adjacent amino acids is approximately 3.5 Å along an extended strand, Cys34, Cys53, and Cys62 are calculated to be 130 Å, 63 Å, and 35 Å distant from the ribosome exit site, respectively. The distributions of PDI and ERp46 molecules bound to RNC 82-aa seem consistent with their accessibility to Cys34 and Cys53, but not to Cys62, as revealed by their mixed disulfide formation with RNC 82-aa (Fig 2D and E).

### Role of the PDI hydrophobic pocket in oxidation of the HSA nascent chain

It is widely known that the PDI **b’** domain contains a hydrophobic pocket that acts as a primary substrate-binding site (Klappa *et al*, 1998). To examine the involvement of the hydrophobic pocket in PDI-catalyzed disulfide bond formation in the HSA nascent chain, we mutated I289, one of the central residues that constitute the hydrophobic pocket, to Ala, and compared the efficiency of disulfide bond introduction into RNC 82-aa between wild-type (WT) and mutant I289A proteins. In this mutant, the x-linker flanked by **b’** and **a’** domains tightly binds the hydrophobic pocket, unlike in WT, thereby preventing PDI from tightly binding an unfolded substrate (Bekendam *et al*, 2016; Nguyen *et al*, 2008). ERp57, another primary member of the PDI family, has a U-shape domain arrangement similar to PDI, but does not contain the hydrophobic pocket in the **b’** domain. For comparison, we also monitored ERp57-catalyzed disulfide introduction into RNC 82-aa.

Despite the occlusion or lack of the hydrophobic substrate-binding pocket, both PDI I289A and ERp57 were found to introduce a disulfide bond into RNC 82-aa at a higher rate than PDI WT (Fig 7A and B). This result suggests that the hydrophobic pocket is involved in binding the HSA nascent chain, but this binding appears to rather slow down disulfide introduction into a nascent chain.

**Figure 7.**
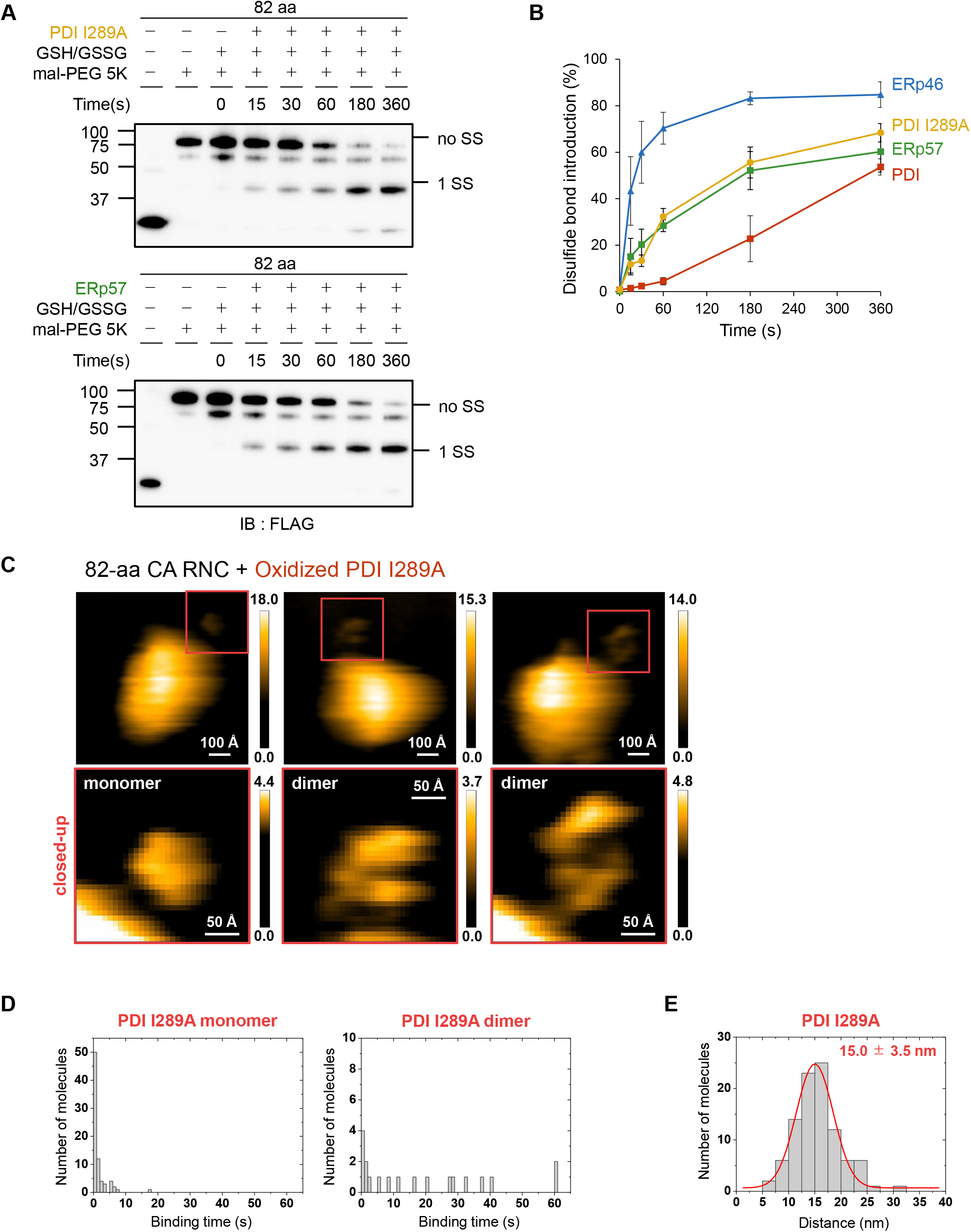
Role of the PDI hydrophobic pocket in PDI-mediated disulfide bond introduction into RNC 82-aa. **A** Disulfide bond introduction into RNC 82-aa by PDI I289A (upper) and ERp57 (lower). Note that faint bands observed between “no SS” and “1SS” likely represent a species in which one of cysteines is not subjected to mal-PEG modification due to glutathionylation. In support of this, these minor bands are even fainter under the conditions of no GSH/GSSG. **B** Quantification of disulfide-bonded species based on the results shown in (A). Quantifications for ERp46 and PDI are based on the results shown in Fig 1E and 1F. n = 3. **C** HS-AFM analyses for binding of PDI I289A to RNC CA 82-aa. Upper AFM images (scan area, 500 Å ’ 500 Å; scale bar, 100 Å) display the PDI I289A molecules that bind 82-aa CA RNC, as marked by red squares. Lower images (scan area, 250 Å × 250 Å; scale bar, 50 Å) highlight the regions surrounded by red squares in the upper images. **D** Histograms show the distribution of the RNC binding time of the PDI I289A monomers (left) and dimers (right). **E** Histogram shows the distribution of the distance between the edge of the ribosome and the centers of RNC-neighboring PDI I289A molecules, calculated from the observed AFM images. Values represents the average distance (mean ± s.d.) calculated from curve fitting with a single-Gaussian model.

To further explore the mechanism by which PDI I289A introduced a disulfide bond at a faster rate than PDI WT, we analyzed its binding to RNC using HS-AFM. The analysis revealed that, while nearly one-third of PDI I289A molecules formed dimers in the presence of RNC 82-aa like PDI WT, the mutant dimers bound RNC for a shorter time than the WT dimers (Fig 7C and Movie EV6). Thus, the RNC-binding time of PDI I289A showed similar distribution to that of ERp46 (Fig 7D and Movies EV5 and EV6), which seems consistent with the higher disulfide introduction efficiency of PDI I289A than that of PDI WT. PDI I289A also bound RNCs at positions ∼16 nm distant from ribosome with a single-Gaussian distribution (Fig 7E), suggesting that PDI I289A recognizes similar sites of the HSA nascent chain as PDI and ERp46.

## Discussion

A number of studies have recently investigated co-translational oxidative folding in the ER (Kadokura *et al*., 2020; Robinson *et al*., 2020; Robinson *et al*., 2017). The present study showed that while both PDI and ERp46 can introduce a disulfide bond into a nascent chain co-translationally, ERp46 catalyzes this reaction more efficiently than PDI and requires a shorter nascent chain segment exposed outside the ribosome exit. Thus, ERp46 appears to be capable of introducing a disulfide bond into a nascent chain during the earlier stages of translation than PDI. The efficient introduction of a Cys53-Cys62 native disulfide on RNC 82-aa by ERp46 (Fig 2) suggests that a separation of ∼8 aa residues between a C-terminal cysteine on a nascent chain and the ribosome exit site (i.e., residues 63-70) is sufficient for ERp46 to catalyze this reaction (Fig 8). When a nascent chain was elongated by the insertion of [SG]-repeat sequences, PDI could also introduce the native disulfide bond into RNCs to some extent (Fig 3B and 3C). Thus, PDI appears to act on a nascent chain to introduce a disulfide bond when the distance between a C-terminal cysteine on a nascent chain and the ribosome exit site reaches ∼18 aa residues (i.e., residues 63-70 + [SG]_5_ repeat; Fig 8).

**Figure 8.**
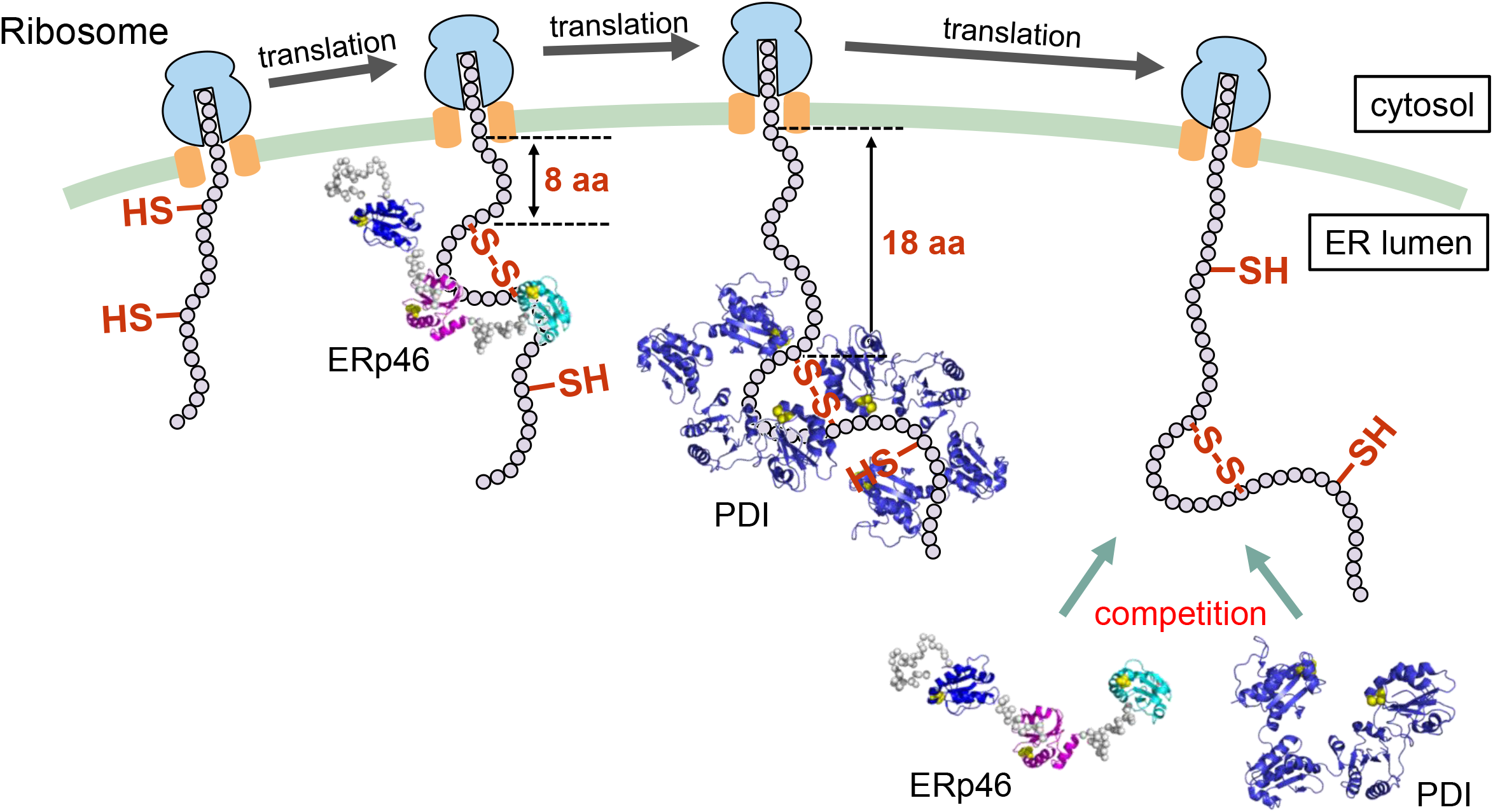
Proposed model of co-translational disulfide bond introduction into nascent chains by ERp46 and PDI. During the early stages of translation, ERp46 introduces disulfide bonds through transient binding to a nascent chain. For efficient disulfide introduction by ERp46, a pair of cysteines must be exposed by at least ∼8 amino acids from the ribosome exit site. By contrast, PDI introduces disulfide bonds by holding a nascent chain inside the central cavity of the PDI homodimer during the later stages of translation, where a pair of cysteines must be exposed by at least ∼18 amino acids from the ribosome exit site. However, when a longer polypeptide is exposed outside the ribosome, ERp46- or PDI-mediated disulfide bond formation can be slower, possibly due to formation of higher-order conformation in the nascent chain. Longer nascent chains may allow PDI family enzymes to compete with each other for binding and acting on RNC.

Disulfide bond formation in partially ER-exposed nascent chains was indeed observed with the ADAM10 disintegrin domain, which has a dense disulfide bonding pattern and little defined structure (Robinson *et al*., 2020). Thus, disulfide bond formation seems to be allowed before the higher order structure is defined in a nascent chain. This could be the case with a Cys34-Cys53 nonnative disulfide and a Cys53-Cys62 native disulfide on RNC 82-aa, since the N-terminal 82-residue HSA fragment alone is unlikely to fold to a globular native-like structure though the fragment of residue 35 to 56 is predicted to form an α-helix according to the HSA native structure. In contrast, some proteins including β2-microglobulin (β2M) and prolactin are shown to form disulfide bonds only after a folding domain is fully exposed to the ER or a polypeptide chain is released from ribosome, suggesting their folding-driven disulfide bond formation. Notably, PDI binds β2M when the N-terminal ∼80 residues of β2M are exposed to the ER, and completes disulfide bond introduction at the even later stages of translation (Robinson *et al*., 2017). Thus, PDI has been demonstrated to engage in disulfide bond formation during late stages of translation or after translation in the ER.

Regarding mechanistic insight, the present HS-AFM analysis visualized PDI and ERp46 acting on nascent chains at the single-molecule level. We found that PDI forms a face-to-face homodimer that binds a nascent chain, as is the case with reduced and denatured full-length substrates (Okumura *et al*., 2019). On the other hand, ERp46 maintains a monomeric form while binding a nascent chain. Interestingly, the PDI dimer binds a nascent chain much more persistently than the PDI monomer and ERp46, suggesting that the PDI dimer holds a nascent chain tightly inside its central hydrophobic cavity. In agreement with this observation, a hydrophobic-pocket mutant (I289A) of PDI bound a nascent chain for shorter time and introduced a disulfide bond into a nascent chain more rapidly than the WT enzyme, as was the case with ERp46. In this context, PDI competed with ERp46 for acting on RNC 95-aa, and thereby inhibited ERp46-mediated disulfide introduction (Fig 4 and Fig 8). Thus, PDI family enzymes do not always work synergistically to accelerate oxidative protein folding, but may possibly inhibit each other during co-translational disulfide bond formation.

How the ER membrane translocon channel is involved in co-translational oxidative folding catalyzed by PDI family enzymes remains an important question. It is possible that PDI and ERp46 form a supramolecular complex with ribosomes and the Sec61 translocon channel via a nascent chain. Indeed, PDI was previously identified as a luminal protein that was in close contact with translocating nascent chains (Klappa *et al*, 1995). Additionally, the oligosaccharyltransferase complex (Harada *et al*, 2009) and an ER chaperone calnexin (Farmery *et al*, 2000) have been reported to interact with the ribosome-associated Sec61 channel to catalyze N-glycosylation and folding of nascent chains in the ER, respectively. In this regard, it will be interesting to examine the close co-localization of PDI/ERp46 with the Sec61 channel in the presence or absence of nascent chains in transit into the ER lumen by super-resolution microscopy or other tools. Systematic studies with a wider range of substrates of different lengths from the ribosome exit site and different numbers of cysteine pairs, and with other PDI family members potentially having different functional roles, will provide further mechanistic and physiological insights into co-translational oxidative folding and protein quality control in the ER.

## Materials & Methods

### Construction of HSA plasmids

DNA fragments encoding specific regions (69-aa, N-terminal pro-sequence 6-aa + the subsequent 63-aa; 82-aa, N-terminal pro-sequence 6-aa + the subsequent 76-aa; 95-aa, N-terminal pro-sequence 6-aa + the subsequent 89-aa) of HSA were amplified by PCR with appropriate primers and inserted into the pUC-T7-HCV-FLAG-2A-uORF expression plasmid, as described in Machida et al. (2014). The amplified fragments were replaced with the 2A region to generate pUC-T7-HCV-FLAG-HSA (69-aa or 82-aa)-uORF2. RNC 82-aa C34A/C53A/C62A and mono-Cys mutants were constructed using the QuikChange method with appropriate primers (Table 1). RNC 82-aa C34A with [SG]_5_ or [SG]_10_ repeats were constructed by the Prime STAR MAX (Takara Bio Inc., Japan) method using appropriate primers (Table 1).

**Table 1.**
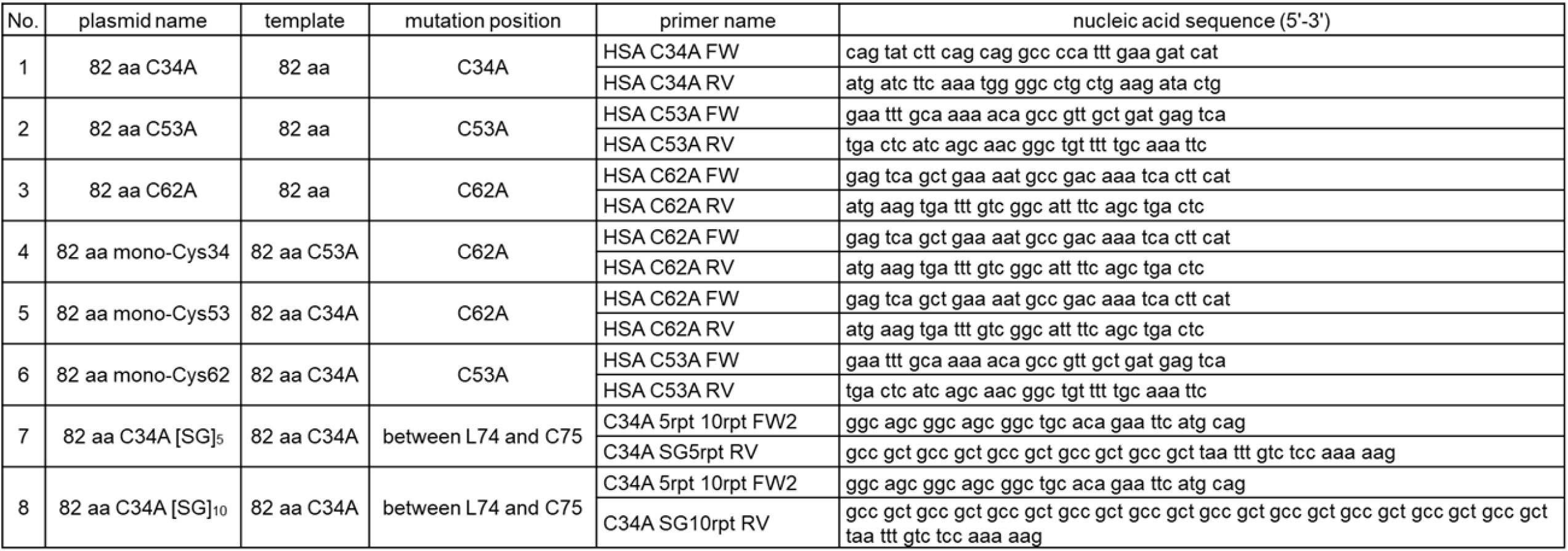
Primers used in this study

### Expression and purification of PDI and ERp46

Overexpression and purification of human PDI and ERp46, and their mutants, were performed as described previously (Kojima *et al*., 2014; Sato *et al*., 2013). An ERp46 trapping mutant with a CXXA sequence in all Trx-like domains was constructed by the QuikChange method using appropriate sets of primers.

### Preparation of RNCs using a translation system reconstituted with human factors

A cell-free translation system was reconstituted with eEF1 (50 μM), eEF2 (1 μM), eRF1/3 (0.5 μM), aminoacyl-tRNA synthetases (0.15 μg/μl), tRNAs (1 μg/μl), 40S ribosomal subunit (0.5 μM), 60S ribosomal subunit (0.5 μM), PPA1 (0.0125 μM), amino acids mixture (0.1 mM) and T7 RNA polymerase (0.015 μg/μl) (Machida *et al*., 2014). We added 1.0 µL template plasmid (0.5 mg/mL) into 19 µL of this cell-free system, and the mixture was incubated for at least 3−4.5 h at 32°C. After HKMS buffer (comprising 25 mM HEPES-KOH (pH 7.0), 150 mM KCl, 5 mM Mg(OAc)_2_, and 1.0 M sucrose) was added, samples were ultra-centrifuged at 100,000 g overnight at 4 °C to recover the RNC as a pellet. After removing the supernatant, pellets were resuspended in HKM buffer comprising 25 mM HEPES-KOH (pH 7.0), 150 mM KCl, and 5 mM Mg(OAc)_2_.

### Monitoring PDI- and ERp46-mediated disulfide bond introduction into RNCs

The RNC suspension prepared as described above was mixed with PDI or ERp46 (0.1 μM each) and glutathione/oxidized glutathione (GSH/GSSG; 1.0 mM:0.2 mM; NACALAI TESQUE, INC., Japan). Aliquots were collected after incubation at 30°C for the indicated times, and reactions were quenched with mal-PEG 5K (2 mM; NOF CORPORATION, Japan) for RNC 69-aa and RNC 82-aa. After cysteine alkylation at room temperature for 20 min, samples were separated by 12% Bis-Tris (pH7.0) PAGE (Thermo Fisher Scientific K.K., Japan) in the presence of the reducing reagent β-mercaptoethanol (β-ME; 10% v/v; NACALAI TESQUE, INC., Japan). After transferring onto a polyvinylidene fluoride (PVDF) membrane (Merck KGaA, Darmstadt, Germany), bands on the membrane were visualized using Chemi-Lumi One Ultra (NACALAI TESQUE, INC., Japan) and a ChemiDocTM Imaging System (Bio-Rad Laboratories, Inc., CA, USA). Signal intensity was quantified using ImageLab software (Bio-Rad Laboratories, Inc., CA, USA).

For RNC 95-aa, reactions were quenched with PEG-PCMal (Dojindo, Japan). After cysteine alkylation at room temperature for 20 min, samples were separated by 10% Bis-Tris (pH7.0) PAGE (Thermo Fisher Scientific K.K., Japan) in the presence of the reducing reagent β-ME (10% v/v;). After gel electrophoresis, the gel was subjected to UV irradiation (302 nm, 8 W) for 30 min. The subsequent procedures were the same as described above.

### Monitoring intermolecular disulfide bond linkage between PDI/ERp46 and ribosome-HSA nascent chain complexes

To detect the intermolecular disulfide bond linkage between PDI/ERp46 and the ribosome-HSA nascent chain complex, we employed RNC 82-aa mono-Cys mutants retaining one of Cys34, Cys53, or Cys62. The RNC suspension prepared as described above was mixed with a PDI or ERp46 trapping mutant (1 μM each) and diamide (100 µM). Aliquots were collected after incubation at 30°C for 10 min, and reactions were quenched with N-ethylmaleimide (2 mM; NACALAI TESQUE, INC., Japan). Samples were analyzed by Nu-PAGE and western blotting as described above.

### High-speed atomic force microscopy imaging

The structural dynamics of PDI and ERp46 were probed using a high-speed AFM instrument developed by Toshio Ando’s group (Kanazawa University). Data acquisition for ERp46 was performed as described previously (Okumura *et al*., 2019). Briefly, His_6_-tagged ERp46 was immobilized on a Co^2+^-coated mica surface through the N-terminal His-tag. To this end, a droplet (10 μL) containing 1 nM ERp46 was loaded onto the mica surface. After a 3 min incubation, the surface was washed with TRIS buffer (50 mM TRIS-HCl pH7.4, 300 mM NaCl). Single-molecule imaging was performed in tapping mode (spring constant, ∼0.1 N/m; resonant frequency, 0.8–1 MHz; quality factor in water, ∼2) and analyzed using Kodec4.4.7.39 software developed by Toshio Ando’s group (Kanazawa University). AFM observations were made in fixed imaging areas (400 × 400 Å^2^) at a scan rate of 0.1 s/frame. Each molecule was observed separately on a single frame with the highest pixel setting (60 × 60 pixels). Cantilevers (Olympus, Tokyo, Japan) were 6–7 μm long, 2 μm wide, and 90 nm thick. For AFM imaging, the free oscillation amplitude was set to ∼1 nm, and the set-point amplitude was around 80% of the free oscillation amplitude. The estimated tapping force was <30 pN. A low-pass filter was used to remove noise from acquired images. The area of a single ERp46 molecule in each frame was calculated using LabView 2013 (National Instruments, Austin, TX, USA) with custom-made programs.

To observe the binding of PDI/ERp46 to RNCs by HS-AFM, RNCs were immobilized on a Ni^2+^-coated mica surface via electrostatic interactions. To this end, a droplet (10 μL) containing RNCs was loaded onto the mica surface. After a 10 min incubation, the surface was washed with HSA buffer comprising 25 mM HEPES-KOH pH 7.0, 150 mM KCl, and 5 mM Mg(OAc)_2_. PDI/ERp46 lacking the N-terminal His_6_-tag was added to the RNC-immobilized mica surface at a final concentration of 1 nM. Measurements were performed under the same conditions described above.

## Acknowledgments

This work was supported by Grants-in-Aid for Scientific Research from MEXT to KI (26116005 and 18H03978), the NAGASE Science Technology Foundation (K.I.) and the MITSUBISHI Foundation (K.I.). This work was also supported by Grant-in-Aid for JSPS Fellows (Grant Number 20J11932 to C.H.) and a Grant-in-Aid of Tohoku University, Division for Interdisciplinary Advanced Research and Education (to C.H.).

## Author contributions

C.H. and T.M. developed an experimental system for directly monitoring co-translational disulfide bond formation. K.M. and H.I. developed and prepared cell-free protein translation system reconstituted with human factors. C.H. prepared various plasmids. C.H. and M.O. purified PDI and ERp46, and their mutants. C.H. and K.N. performed HS-AFM measurements and analyses. C.H., K.N., M.O. and T.O. discussed the results of HS-AFM. K.I. supervised the work. C.H. and K.N. prepared the Figures. C.H. and K.I. wrote the manuscript. All of the authors discussed the results and approved the manuscript.

## Expanded View

**Figure EV1.**
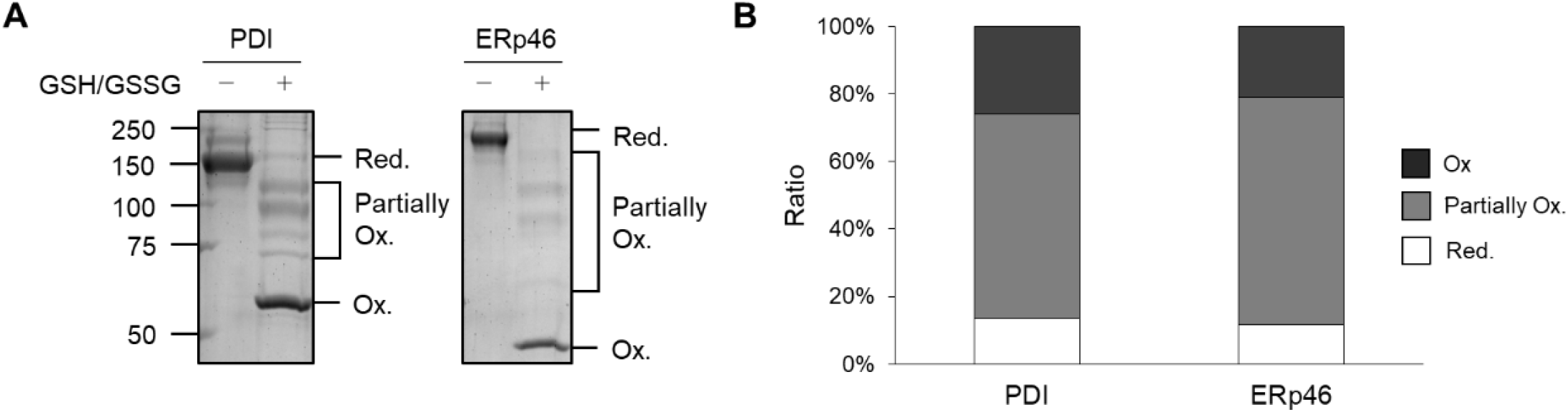
Redox states of PDI and ERp46 in glutathione redox buffer and disulfide bond introduction into 82 aa C34A, catalyzed by PDI a domain. **A** Redox states of PDI and ERp46 in the presence of 1 mM GSH and 0.2 mM GSSG. Purified PDI and ERp46 were incubated for 6 mins at 30 °C in the above glutathione redox buffer and modified with 2 mM mal-PEG 5K for separation on SDS gels. **B** Quantification based on the results shown in (A).

**Figure EV2.**
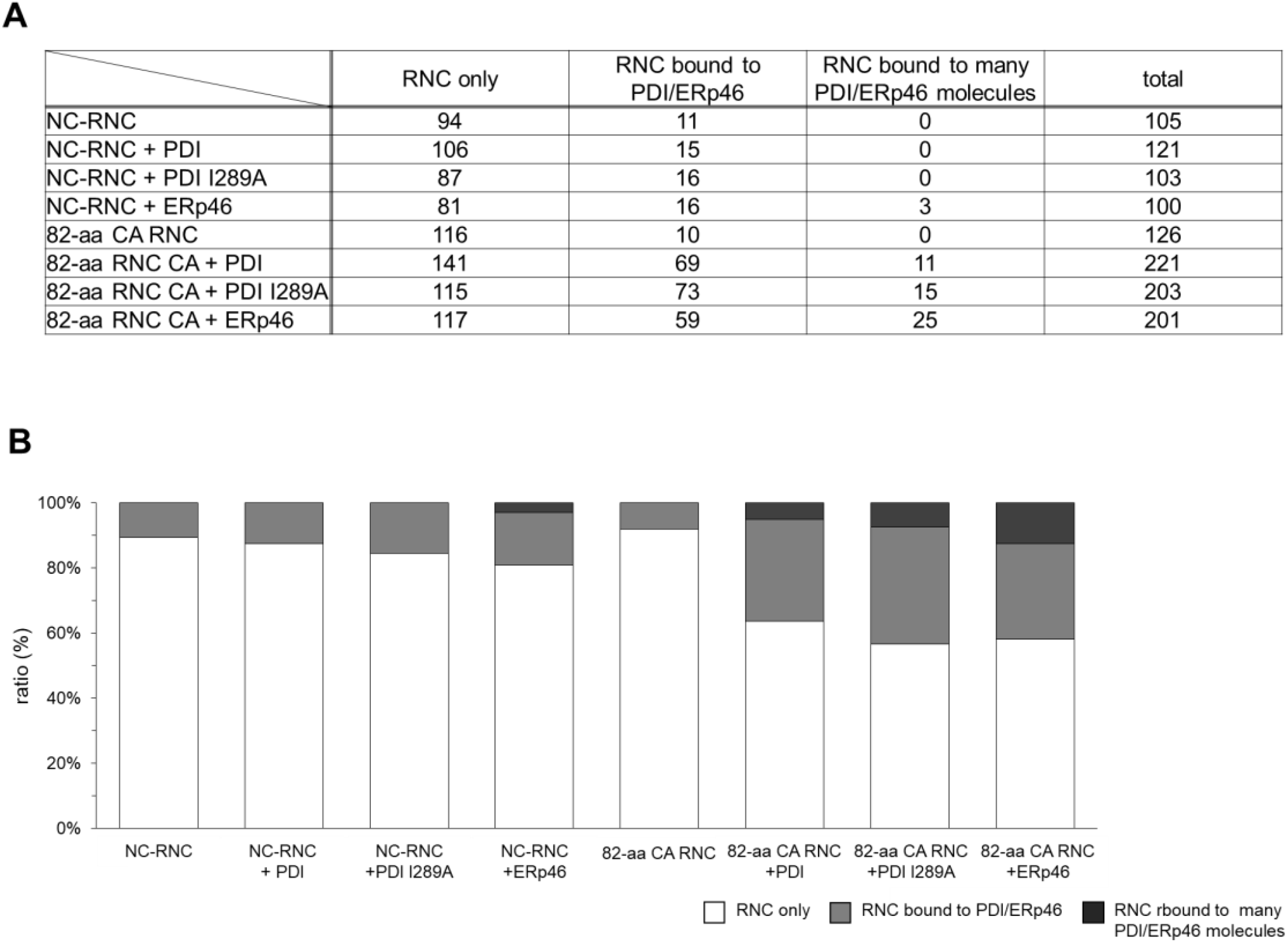
Statistical analysis of RNC molecules observed by HS-AFM in the presence or absence of PDI/ERp46. **A** Number of particles observed for NC-RNC or 82-aa CA RNC molecules present in isolation or bound to PDI/ERp46 molecules. **B** Ratio of NC-RNC or 82-aa CA RNC molecules present in isolation or bound to PDI/ERp46, calculated based on the observed number of particles in (A). Note that a minor portion of NC-RNC or 82-aa CA RNC molecules were bound to many ERp46/PDI molecules, possibly due to serious structural damages of the RNC molecules.

**Figure EV3.**
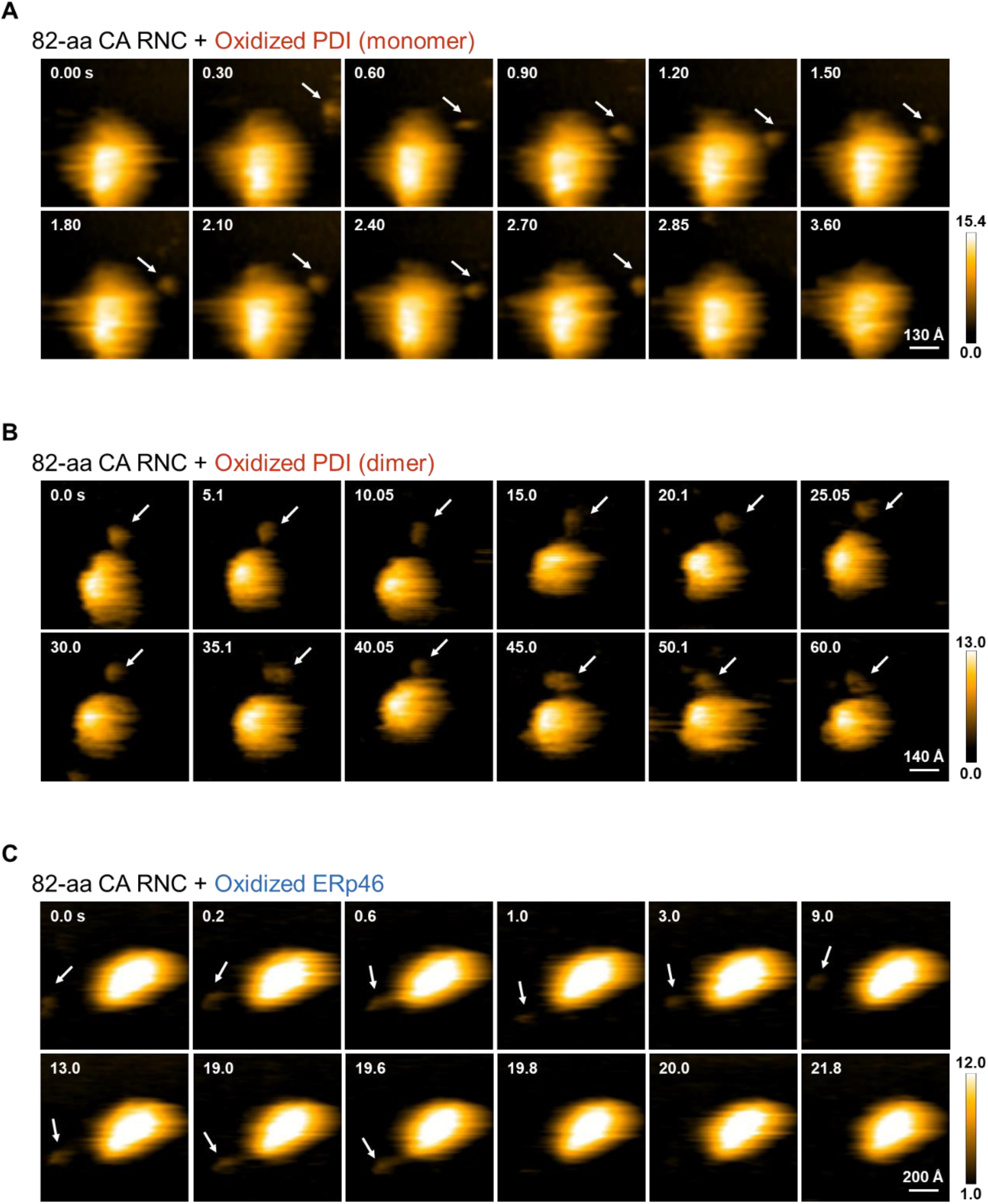
Representative time-course snapshots captured by HS-AFM for 82-aa CA RNC bound to the PDI monomer (A), the PDI dimer (B), and ERp46 (C). **A** Time-course snapshots captured by HS-AFM for the PDI monomer binding to 82-aa CA RNC. The AFM images (scan area, 650 Å ’ 650 Å; scale bar, 130 Å) displaying 82-aa CA RNC in the presence of oxidized PDI (1 µM). White arrows indicate the monomeric PDI molecules that bind to 82-aa CA RNC. See also supplementary video 2. **B** Time-course snapshots captured by HS-AFM for the PDI dimer binding to 82-aa CA RNC. The AFM images (scan area, 700 Å ’ 700 Å; scale bar, 140 Å) displaying 82-aa CA RNC in the presence of oxidized PDI (1 µM). White arrows indicate the dimeric PDI molecules that bind to 82-aa CA RNC. See also supplementary video 3. **C** Time-course snapshots captured by HS-AFM for ERp46 binding to 82-aa CA RNC. The AFM images (scan area, 1,000 Å ’ 1,000 Å; scale bar, 200 Å) displaying 82-aa CA RNC in the presence of oxidized ERp46 (1 µM). White arrows indicate the ERp46 molecules that bind to 82-aa CA RNC. See also supplementary video 4.

**Figure EV4.**
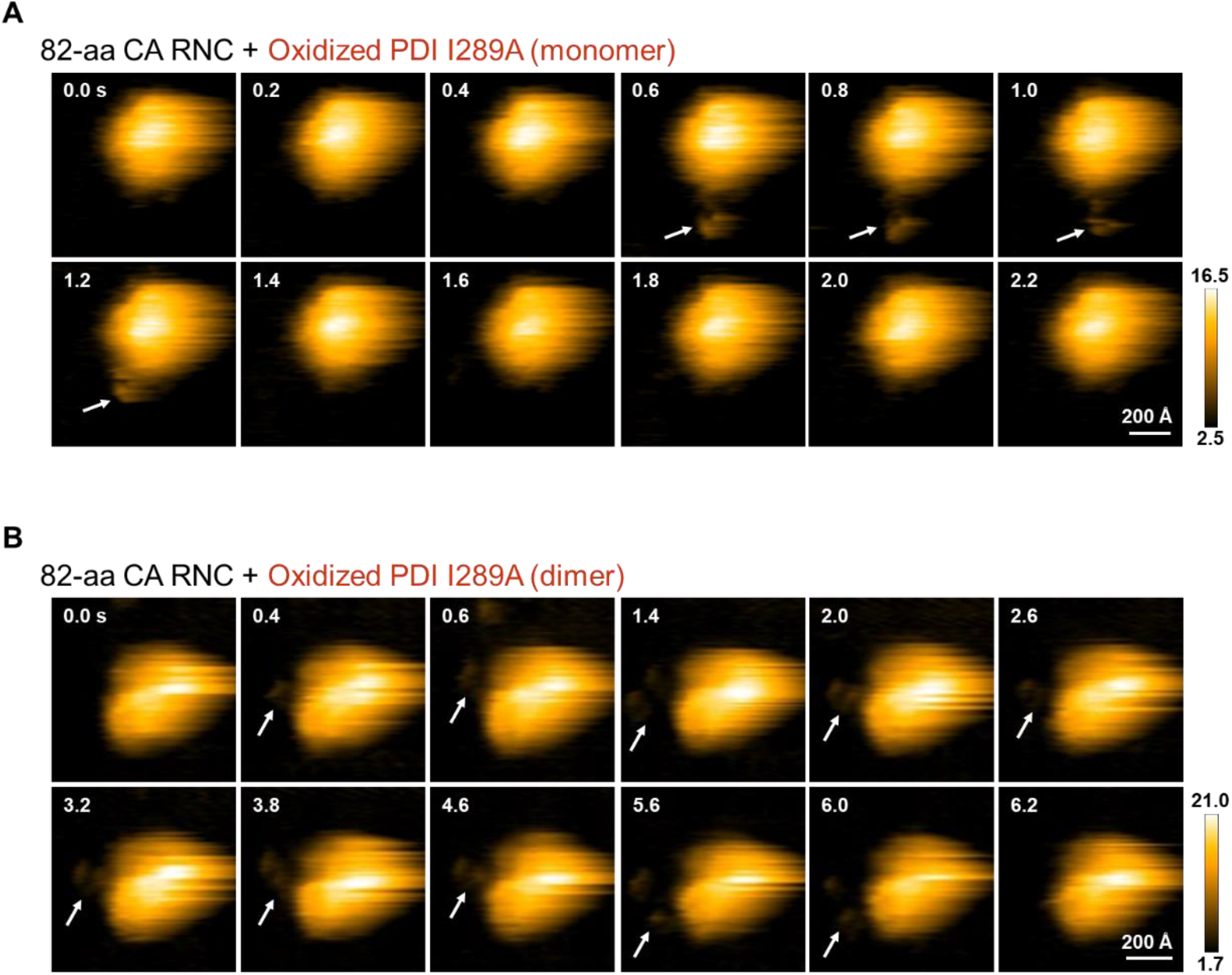
Representative time-course snapshots captured by HS-AFM for 82-aa CA RNC bound to the PDI I289A monomer (A), and the PDI I289A dimer (B). **A** Time-course snapshots captured by HS-AFM for the PDI I289A monomer binding to 82-aa CA RNC. The AFM images (scan area, 900 Å ’ 900 Å; scale bar, 200 Å) displaying 82-aa CA RNC in the presence of oxidized PDI I289A (1 µM). White arrows indicate the monomeric PDI I289A molecules that bind to 82-aa CA RNC. See also supplementary video 5. **B** Time-course snapshots captured by HS-AFM for the PDI I289A dimer binding to 82-aa CA RNC. The AFM images (scan area, 800 Å ’ 800 Å; scale bar, 200 Å) displaying 82-aa CA RNC in the presence of oxidized PDI I289A (1 µM). White arrows indicate the dimeric PDI I289A molecules that bind to 82-aa CA RNC. See also supplementary video 6.

**Movie EV1 - HS-AFM movies showing structure dynamics of oxidized ERp46**. This movie is a source of the time-course snapshots shown in Fig 5C.

**Movie EV2 - HS-AFM movies showing the binding of the PDI monomer to 82-aa CA RNC**. This movie is a source of the time-course snapshots shown in supplementary Fig EV3A.

**Movie EV3 - HS-AFM movies showing the binding of the PDI dimer to 82-aa CA RNC**. This movie is a source of the time-course snapshots shown in supplementary Fig EV3B.

**Movie EV4 - HS-AFM movies showing the binding of ERp46 to 82-aa CA RNC**. This movie is a source of the time-course snapshots shown in supplementary Fig EV3C.

**Movie EV5 - HS-AFM movies showing the binding of the PDI I289A monomer to 82-aa CA RNC**. This movie is a source of the time-course snapshots shown in supplementary Fig EV4A.

**Movie EV6 - HS-AFM movies showing the binding of the PDI I289A dimer to 82-aa CA RNC**. This movie is a source of the time-course snapshots shown in supplementary Fig EV4B.

